# *β*-catenin-Mediated Wnt Signal Transduction Proceeds Through an Endocytosis-Independent Mechanism

**DOI:** 10.1101/2020.02.13.948380

**Authors:** Ellen Youngsoo Rim, Leigh Katherine Kinney, Roel Nusse

## Abstract

The Wnt pathway is a key intercellular signaling cascade that regulates development, tissue homeostasis, and regeneration. However, gaps remain in our understanding of the molecular events that take place between ligand-receptor binding and target gene transcription. Here we used a novel tool for quantitative, real-time assessment of endogenous pathway activation, measured in single cells, to answer an unresolved question in the field – whether receptor endocytosis is required for Wnt signal transduction. We combined knockdown or knockout of essential components of Clathrin-mediated endocytosis with quantitative assessment of Wnt signal transduction in mouse embryonic stem cells (mESCs). Disruption of Clathrin-mediated endocytosis did not affect accumulation and nuclear translocation of *β*-catenin, as measured by single-cell live imaging of endogenous *β*-catenin, and subsequent target gene transcription. Disruption of another receptor endocytosis pathway, Caveolin-mediated endocytosis, did not affect Wnt pathway activation either. These results, confirmed in multiple cell lines, suggest that endocytosis is not a general requirement for Wnt signal transduction. We show that off-target effects of a drug used to inhibit endocytosis may be one source of the discrepancy among reports on the role of endocytosis in Wnt signaling.

## Introduction

The Wnt pathway is an intercellular signaling pathway activated by secreted glycoproteins of the Wnt family. Conserved in all multicellular animals, Wnt pathway activity is important for developmental decisions such as embryonic axis establishment and cell fate determination (1, 2). In the adult organism, Wnt activity is essential for maintenance of stem cells in many tissues, including the intestine, the skin, and the blood (3–5). Hyperactivation of the Wnt pathway, on the other hand, is implicated in human disease including multiple types of cancer (6, 7). Therefore, precise regulation of Wnt signaling is essential for normal development and proper maintenance of adult tissues.

There are branches of the Wnt signaling pathway that utilize overlapping sets of molecular effectors to elicit different biological outcomes. Of these, the *β*-catenin-dependent branch of Wnt signaling is implicated in regulation of cell proliferation and fate specification, and plays an indispensable role in development as well as adult tissue homeostasis and regeneration. In the absence of Wnt ligand-receptor interaction, the signal transducer *β*-catenin is constantly phosphorylated and degraded by GSK3*β* as part of a protein complex. Upon Wnt ligand engagement with co-receptors Frizzled and Lrp5/6 at the cell surface, *β*-catenin phosphorylation by GSK3*β* is inhibited. This leads to *β*-catenin accumulation in the cytoplasm and concomitant translocation into the nucleus, where it can induce transcription of target genes. The importance of *β*-catenin stabilization in Wnt signal transduction has been demonstrated in many *in vivo* and *in vitro* contexts (8, 9). However, immediate molecular responses to the ligand-receptor interaction and how they elicit accumulation of *β*-catenin are not fully elucidated.

One point of uncertainty is whether receptor endocytosis following Wnt binding is required for signal transduction. Endocytosis and endocytic trafficking have been implicated in various signaling pathways. Downregulation of membrane receptors via endocytosis dampens further signaling in some contexts whereas internalization of receptor complexes enhances signal transduction in others (10). Two endocytic pathways, Caveolin-mediated and Clathrin-mediated endocytosis, have been implicated in receptor internalization in Wnt signaling (11–16). In Caveolin-mediated endocytosis, Caveolin polymers coat the part of the plasma membrane that undergoes endocytosis. Clathrin-mediated endocytosis, the major endocytic pathway for cellular cargo uptake, initiates when the scaffold protein Clathrin forms a coat at the membrane. Along with the AP2 complex and other adaptor proteins, Clathrin bends the membrane to form a pit, which is pinched off at the neck to form a vesicle that internalizes the cargo (17, 18).

Both Clathrin-and Caveolin-mediated forms of receptor endocytosis have been implicated as essential steps in Wnt signal transduction (11–15). In one model of pathway activation, for instance, internalization of the Wnt receptor via endocytosis activates signaling through formation of multivesicular bodies (MVBs); upon maturation of endocytosed vesicles into MVBs, sequestration of the protein complex including GSK3*β* could allow *β*-catenin to escape degradation and induce target gene activation (19). Other reports suggest that endocytosis negatively regulates Wnt signaling through internalization of receptors, which reduces the level of receptors available for ligand binding at the cell surface (15, 16). One potential mechanism for endocytosis-mediated downregulation of Wnt receptors involves transmembrane ubiquitin ligases RNF43 and ZNRF3. RNF43 and ZNRF3 ubiquitinate Frizzled receptors to induce their endocytic uptake and degradation (20). Inhibition of endocytosis would prevent RNF43/ZNRF3-induced internalization of Frizzled, leading to increased cellular sensitivity to the Wnt signal. Therefore, the field disagrees on 1) whether receptor endocytosis is a positive or negative regulator of Wnt signal transduction, 2) how endocytosis might regulate Wnt signaling in each case, and 3) which endocytic pathway and molecular effectors might be involved.

These questions call for a careful examination of the role of receptor endocytosis in Wnt signaling. To investigate this, we employed knockdown or CRISPR-mediated knockout of essential endocytic components then assessed their effect on Wnt signal transduction. We used multiple cell types, including mouse embryonic stem cells (mESCs), which are inherently Wnt-responsive (21, 22). As we show, mESCs can be used at the single-cell level free of other cell types that might confound the effect of endocytosis on signal transduction in Wnt receiving cells. To monitor activation of the native Wnt pathway in these cells, we used readouts such as live imaging nuclear accumulation of endogenous *β*-catenin and subsequent target gene transcription. Our results in mESCs and additional cell lines demonstrate that Clathrin-mediated endocytosis is not required for *β*-catenin stabilization and target transcription. Similarly, disruption of Caveolin-mediated endocytosis does not affect Wnt pathway activation.

## Results

### *β*-catenin Knock-in Reporter Provides a Faithful Read-out for Wnt Activity in mESCs

mESCs are equipped with the molecular machinery for Wnt signal transduction and depend on Wnt signaling for self-renewal and maintenance of pluripotency (21, 23, 24), presenting a functionally relevant context to study the potential role of endocytosis in Wnt signaling. To determine whether receptor endocytosis is required for Wnt activation, we needed to quantitatively evaluate Wnt signal transduction in mESCs in which endocytic function was disrupted. We sought to rely on native Wnt pathway components to assess cellular responses to Wnt stimulation. Therefore, we used a mESC knock-in reporter line to track endogenous *β*-catenin in single cells. The cell line harbors a functional allele of *β*-catenin fused at the C-terminus to the yellow fluorescent protein Venus. To visualize the cell nucleus and track nuclear *β*-catenin over time, cyan fluorescent protein mTurquoise-tagged H2B was introduced into this reporter cell line (Fig 1A).

**Fig. 1.**
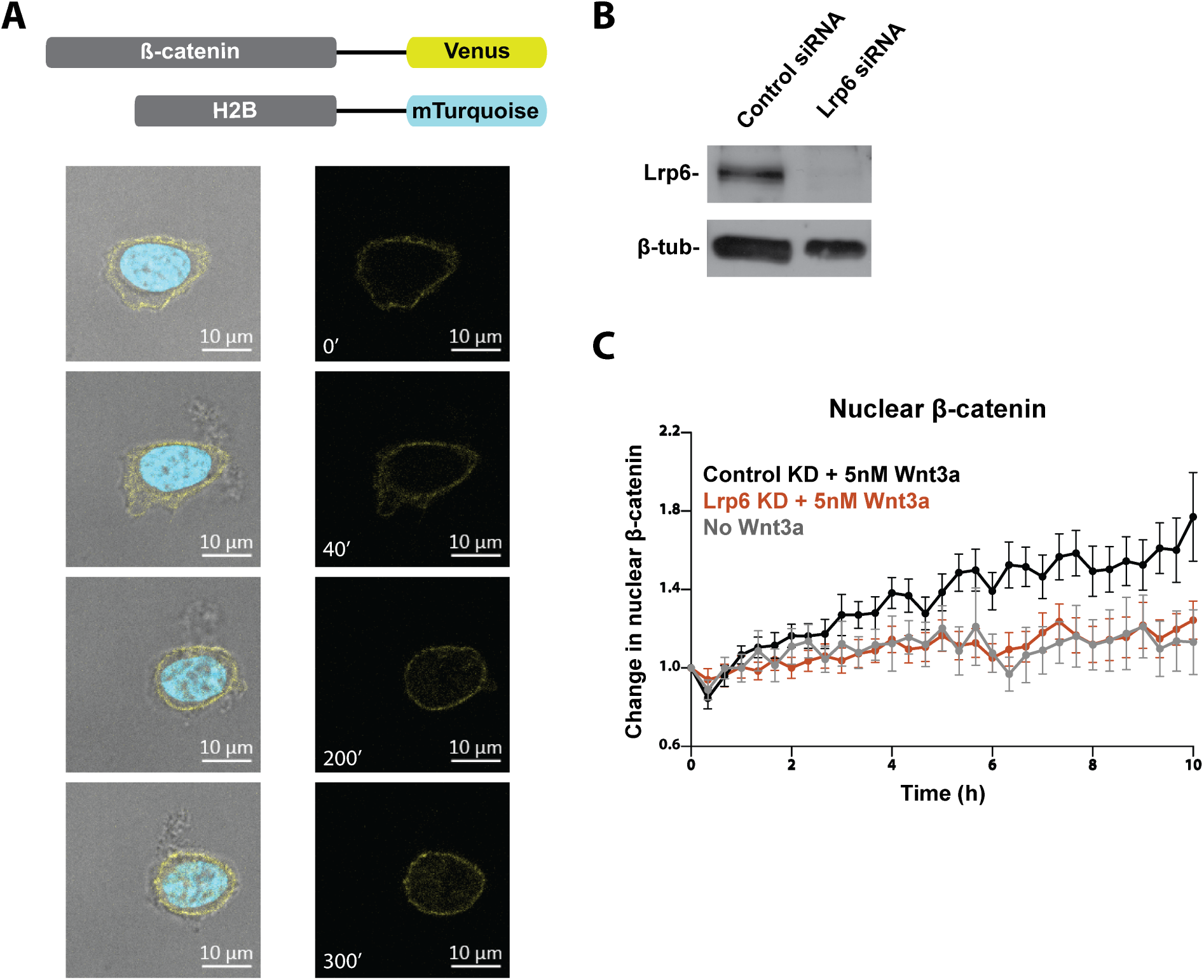
A *β*-catenin knock-in reporter provides a faithful readout for Wnt activity in mESCs. (A) Endogenous *β*-catenin was tagged at the C-terminus with Venus and the histone protein H2B was tagged at the C-terminus with mTurquoise. Snapshots of a mESC harboring the *β*-catenin knock-in reporter show gradual increase in the amount of *β*-catenin (yellow) in the nucleus (cyan). Images were taken over 5 hours following stimulation with 5nM Wnt3a. Scale bar, 10*μ*m. (B) Effective siRNA-mediated knockdown of Lrp6 was confirmed via Western blot. (C) Change in nuclear *β*-catenin upon stimulation with 5nM Wnt3a was quantified. Lrp6 knockdown ablated nuclear translocation of *β*-catenin. Single *β*-catenin reporter mESCs were live imaged over 10 hours. (Control siRNA, n=23; Lrp6 siRNA, n=22; no Wnt3a, n=16).

In mESC colonies, the fusion protein reported *β*-catenin localization at the cell membrane, while addition of Wnt3a protein induced its nuclear translocation (Fig 1A). Single cells live imaged for up to 10 hours were used to quantify cellular accumulation and nuclear translocation of *β*-catenin. Consistent with previous reports of *β*-catenin dynamics upon pathway activation, individual cells presented significant variability in their response in terms of the rate and extent of change in *β*-catenin (8, 25, 26). Some cells exhibited substantial increase in total and nuclear *β*-catenin within an hour while others showed slow or minor increase, reflecting the inherent variability in the cells’ molecular machinery to respond to Wnt signaling. However, when *β*-catenin dynamics were quantified and averaged in 20 to 30 single cells, Wnt3a treatment consistently led to 1.5 to 2-fold increase in the nuclear fraction of *β*-catenin (Fig 1C and Fig 2C) and 2 to 3-fold increase in overall cellular *β*-catenin (Fig 2D). Nuclear *β*-catenin manifested in a speckled pattern, possibly reflecting incorporation into large nuclear complexes.

**Fig. 2.**
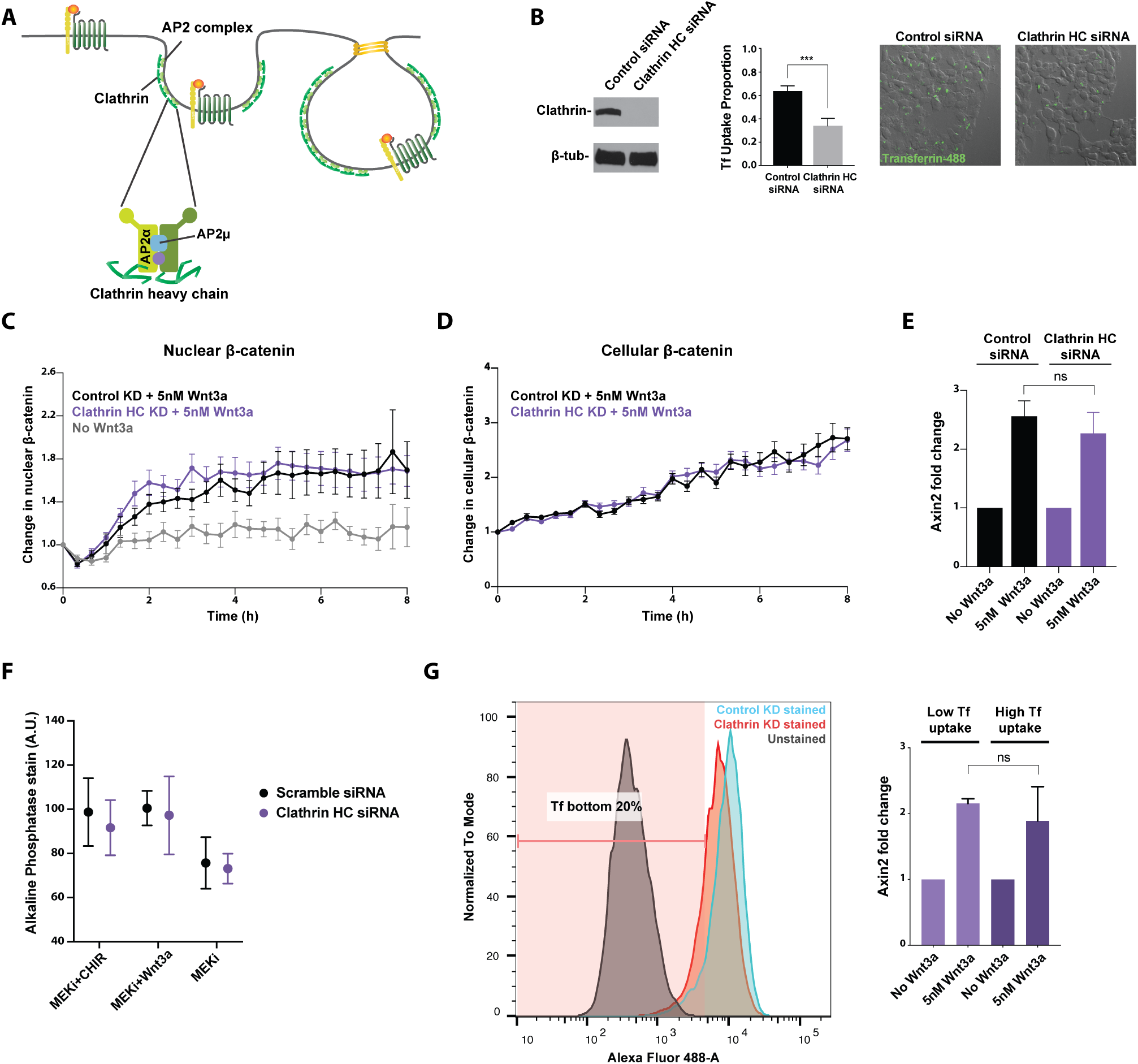
Knockdown of Clathrin heavy chain impairs Clathrin-mediated cargo uptake but does not affect Wnt signal transduction. (A) Clathrin and the AP2 complex are key components of the Clathrin endocytic pathway. Clathrin HC and AP2 subunits *α* and *μ* were targeted to inhibit Clathrin-mediated endocytosis. (B) Effective siRNAmediated knockdown of Clathrin HC was confirmed via Western blot. Clathrin HC knockdown impaired cellular uptake of fluorescently labeled Transferrin as demonstrated by a significant reduction in the proportion of cells with Transferrin uptake (p *<* 0.001). (C) Change in nuclear and cellular *β*-catenin upon stimulation with 5nM Wnt3a was quantified. Clathrin HC knockdown did not affect nuclear translocation or cellular accumulation of *β*-catenin. (Control siRNA, n=22; Clathrin HC siRNA, n=22; no Wnt3a, n=15). (D) Transcription of the Wnt target gene Axin2 was unaffected by knockdown of Clathrin HC. (E) Alkaline phosphatase stain intensity was quantified for each colony in the specified culture condition (Control siRNA MEKi+CHIR, n=24; Clathrin HC siRNA MEKi+CHIR, n=12; Control siRNA MEKi+Wnt3a, n=29; Clathrin HC siRNA MEKi+Wnt3a, n=22; Control siRNA MEKi, n=23; Clathrin HC siRNA MEKi, n=21). (F) Cells in the bottom 20% of Transferrin uptake function, as measured by low intracellular fluorescent Transferrin signal, were sorted via FACS. Axin2 transcription was induced in these cells to a level similar to cells in the top 80% of Transferrin uptake.

As expected, knockdown of the essential Wnt receptor Lrp6 led to loss of *β*-catenin accumulation in the nucleus even in the presence of Wnt3a (Fig 1B and C). Therefore, the reporter mESCs allow visual assessment of Wnt pathway activation in a quantitative, temporally sensitive manner, and can be used to study potential effects of disruption in endocytosis on Wnt signal transduction.

### Knockdown of Clathrin Heavy Chain Does Not Affect Wnt Signal Transduction

The scaffold protein Clathrin forms a coat at the membrane to initiate Clathrin-mediated endocytosis (17). The AP2 complex and other adaptor proteins aid Clathrin in bending the membrane to form a pit, which is pinched off and internalized as a vesicle (Fig 2A). To determine whether Clathrin-mediated endocytosis is required for Wnt activation, we first targeted the main structural component of the Clathrin endocytic pit, Clathrin heavy chain (HC), via siRNA-mediated knockdown. To ensure that Clathrin-mediated endocytosis was impaired in these cells, we assessed cellular uptake of Transferrin, a major cargo of Clathrin endocytosis (27). Significant reduction in the uptake of fluophore-conjugated Transferrin was observed in cells with Clathrin HC knockdown (Fig 2B).

Surprisingly, reporter mESCs with impaired Clathrin endocytosis exhibited similar kinetics in both cellular accumulation and nuclear translocation of *β*-catenin (Fig 2C and 2D). Consistent with appropriate Wnt pathway activation, transcription of Axin2 was induced in mESCs subjected to Clathrin HC knockdown just as in cells subjected to mock knockdown (Fig 2E).

Wnt activation and consequent maintenance of pluripotent identity in mESC culture can be achieved through addition of either the GSK3*β* inhibitor CHIR99021 or exogenous Wnt3a (21) (22). Since GSK3*β* inhibition is downstream of Wnt-induced changes at the receptor level, CHIR99021 administration would bypass any requirement for receptor endocytosis in Wnt pathway activation. In contrast, if Clathrin-mediated endocytosis were required for Wnt activation, Clathrin HC knockdown would impair the cells’ ability to maintain pluripotency with exogenous Wnt3a in culture.

mESCs subjected to Clathrin HC knockdown were cultured for three days with CHIR99021, Wnt3a, or neither. We then quantified colony expression of alkaline phosphatase (AP), a hallmark of embryonic stem cell pluripotency (28). In both control and Clathrin HC knockdown conditions, absence of CHIR99021 or Wnt3a in culture led to reduced AP expression as mESCs began to lose pluripotency, whereas CHIR99021 addition prevented loss of AP expression (Fig 2F). Importantly, mESCs subjected to Clathrin HC knockdown were able to maintain AP expression with Wnt3a. These results indicated that impaired Clathrin-mediated endocytosis does not affect maintenance of pluripotency, the key biological consequence of Wnt pathway activation in mESCs.

One concern with the Clathrin HC knockdown experiments was potential retention of Clathrin function due to incomplete knockdown. To address this, we took two approaches: 1) assessing Wnt pathway activation in cells selected for most severe impairment in Clathrin-mediated endocytosis upon Clathrin HC knockdown and 2) knocking out another essential component of the Clathrin pathway. Due to Clathrin’s role in mitotic spindle organization unrelated to its function in endocytosis, knocking out Clathrin HC was not a viable option (29) (30).

First, mESCs were subjected to the Transferrin uptake assay following three-day knockdown of Clathrin HC. Of these, cells in the bottom 20% of Transferrin uptake function were selected via FACS (Fig 2G). When cells in the bottom 20% in terms of Clathrin endocytic function were compared to those in the top 80% and those subjected to mock knockdown, Axin2 was induced to comparable levels (Fig 2G). Therefore, even mESCs most severely compromised in Clathrin endocytosis function retained the ability to respond to Wnt3a stimulation, further supporting that Clathrin-mediated endocytosis is not required for Wnt signal transduction.

### Loss of Adaptor Protein AP2*α* Does Not Affect Wnt Signal Transduction

In a complementary approach to inhibit Clathrin-mediated endocytosis, Ap2*α* was knocked out via CRISPR targeting (Fig 2A). Ap2*α* is one of the large subunits of the heterotetrameric adaptor complex AP2. As AP2 initiates formation of the Clathrin pit on the plasma membrane and mediates the interaction between Clathrin and its cargo proteins, it is an essential component of the Clathrin-mediated endocytic pathway (17).

mESCs were transfected with a CRISPR gRNA targeting exon 4 of Ap2*α* and subjected to clonal selection. An Ap2*α* knockout clone with frameshift deletion alleles was identified (Supplementary Fig 7A) and Western blot confirmed loss of Ap2*α* protein (Fig 3A). Cellular uptake of the Clathrin cargo Transferrin was significantly impaired in mESCs lacking Ap2*α* (Fig 3A). Upon Wnt exposure, cells missing Ap2*α* were capable of accumulating *β*-catenin in the cytoplasm and the nucleus in response to Wnt3a stimulation (Fig 3B and 3C). Transcription of the Wnt target gene Axin2 was properly induced by Wnt3a in the knockout mESCs (Fig 3D). These results are consistent with outcomes of Clathrin HC knockdown and provide further support that Clathrin-mediated endocytosis is dispensable for Wnt signal transduction.

**Fig. 3.**
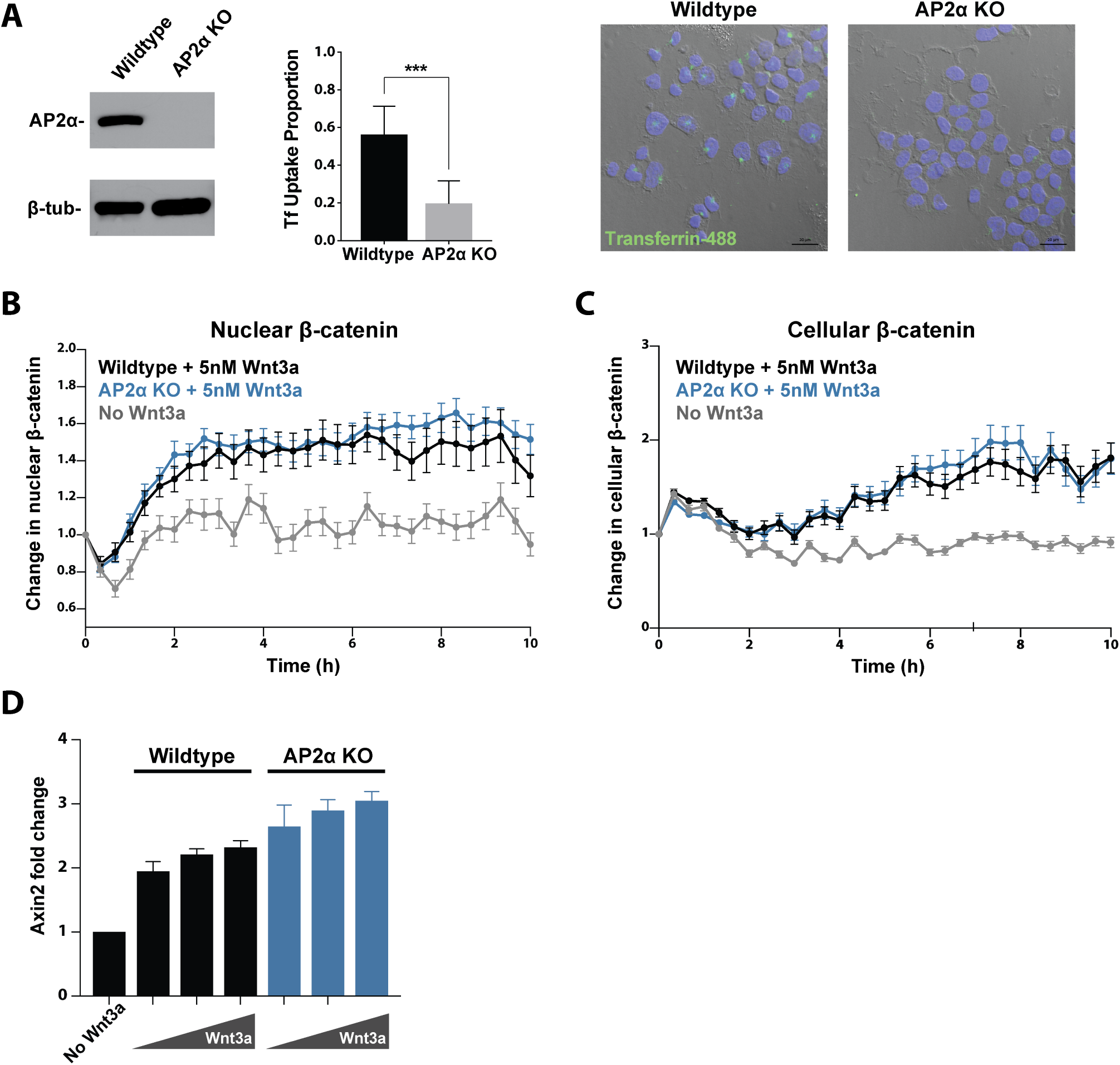
Loss of adaptor protein AP2*α* impairs Clathrin-mediated cargo uptake but does not affect Wnt signal transduction. (A) CRISPR-mediated knockout of AP2*α* was confirmed via Western blot. Loss of AP2*α* impaired cellular uptake of fluorescently labeled Transferrin. (p *<* 0.001). (B) Loss of AP2*α* did not affect nuclear translocation or cellular accumulation of *β*-catenin upon stimulation with 5nM Wnt3a. (Wildtype, n=28; AP2*α* knockout, n=23; no Wnt3a, n=21). (C) Transcription of the Wnt target gene Axin2 upon stimulation with 1nM-5nM Wnt3a was unaffected by loss of AP2*α*.

As mentioned previously, individual cells have been shown to present significant variability in absolute level changes in *β*-catenin (8, 25, 26). Indeed, we observed variability in absolute levels and dynamics of *β*-catenin accumulation among both wildtype and Ap2*α* knockout cells (Supplementary Fig 6A). However, no significant difference in absolute levels of nuclear *β*-catenin on average was observed upon disruption of endocytosis (Supplementary Fig 6B). Therefore, cells lacking Ap2*α* exhibited appropriate relative and absolute increase in nuclear *β*-catenin levels in response to Wnt3a.

### Knockdown of Additional Endocytosis Components Does Not Affect Wnt Signal Transduction

We also disrupted the Clathrin endocytic pathway through knockdown of Ap2m1, which encodes the Ap2*μ* subunit of the AP2 heterotetramer (Fig 2A). Ap2*μ* phosphorylation and subsequent conformation change are necessary for maturation of Clathrin-coated pits (31). Knockdown of Ap2*μ* led to severely impaired Transferrin cargo uptake (Supplementary Fig 7B). Consistent with other disruptions in the Clathrin endocytic pathway, Ap2*μ* knockdown did not affect cytoplasmic accumulation or nuclear translocation of *β*-catenin or Axin2 transcription (Supplementary Fig 7C and 7D).

Some reports in the field have implicated Caveolin-mediated endocytosis in Wnt signal transduction (11) (15). Caveolin expression was barely detectable in the mESC line at the transcript and protein levels (32). Any residual Caveolin function was suppressed with double knockdown of Caveolin 1 and Caveolin 2, two Caveolin proteins that are utilized in non-muscle cell types (33) (Supplementary Fig 8A). Knockdown of both Caveolins did not compromise *β*-catenin accumulation in the nucleus or Axin2 transcription (Supplementary Fig 8B and 8D). Even when knockdown of the Caveolins was combined with knockdown of Clathrin HC, nuclear accumulation of *β*-catenin and consequent Axin2 transcription were unaffected (Supplementary Fig 8C and 8D). These results indicate that neither Clathrin-mediated nor Caveolin-mediated endocytic pathways are required for mESC response to Wnt stimulation.

### Clathrin-mediated Endocytosis Is Not Required for Wnt Signal Transduction in Additional Cell Lines

Given that neither Clathrin- nor Caveolin-mediated endocytic pathways are required for mESC response to Wnt stimulation, we asked whether endocytosis is dispensable for Wnt signal transduction in other cell types. We tested Wnt signal transduction in additional human and mouse cell lines with impaired Clathrin-mediated endocytosis. Clathrin HC was knocked down in LS/L cells which are mouse fibroblasts stably harboring TOPFlash, a luciferase reporter driven by Wnt-responsive elements (13). Induction of Wnt reporter activity by increasing doses of Wnt3a was indistinguishable between LS/L cells subjected to control or Clathrin HC knockdown (Fig 4A). Similarly, knockdown of Clathrin HC in human embryonic kidney 293T cells did not affect their ability to transcribe Axin2 upon Wnt3a stimulation (Fig 4B). Results in these additional cell types support that Clathrin-mediated endocytosis is not a general requirement for Wnt pathway activation.

**Fig. 4.**
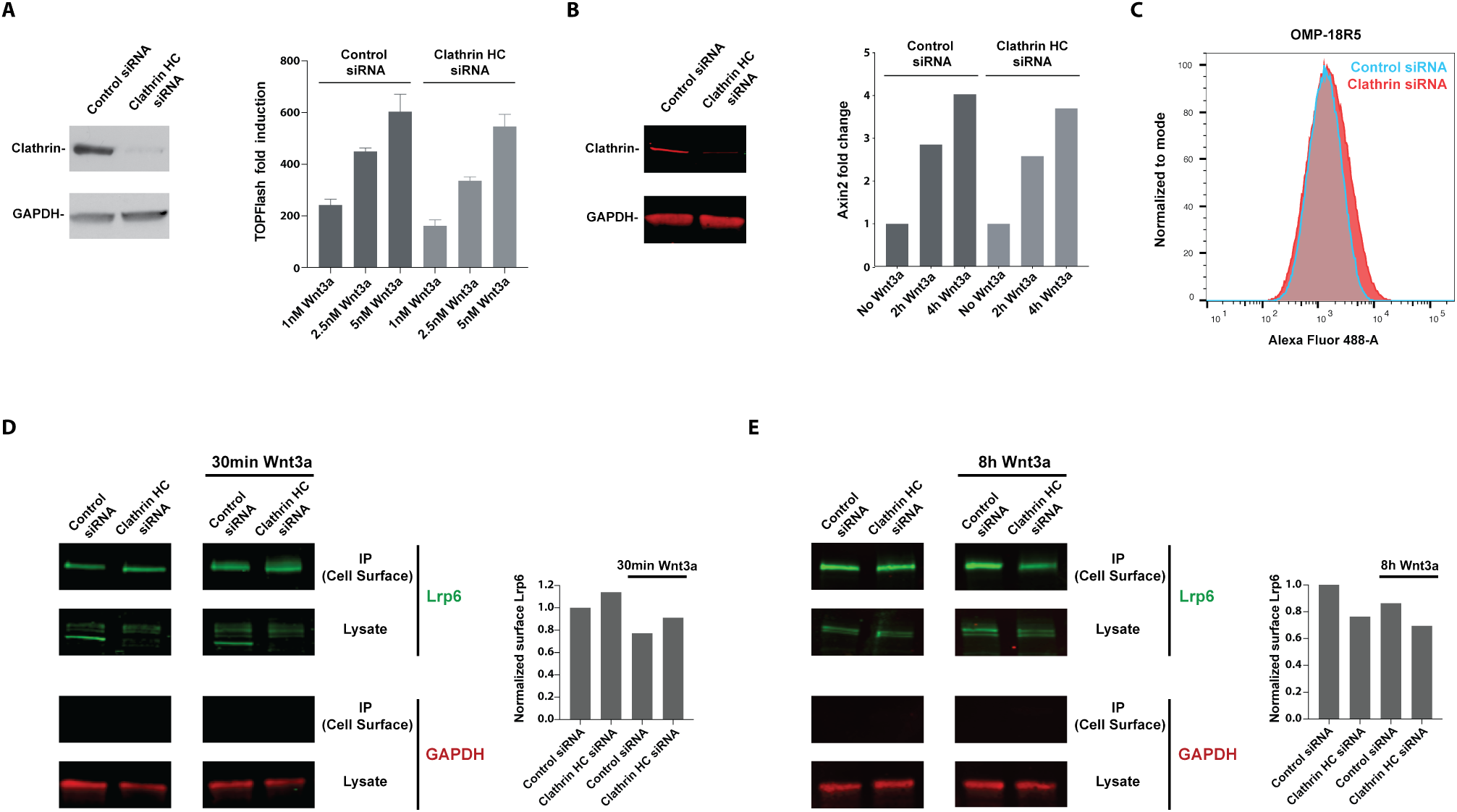
Clathrin-mediated endocytosis is not a significant modulator of Lrp6 or Frizzled receptor abundance on the plasma membrane. (A) Effective knockdown of Clathrin HC in mouse L cells was confirmed via Western blot. Wnt pathway activation, as measured by TOPFlash luciferase activity, by 1nM-5nM Wnt3a was indistinguishable between cells subjected to control or Clathrin HC knockdown. (B) Effective knockdown of Clathrin HC in HEK293T cells was confirmed via Western blot. Transcription of the Wnt target gene Axin2 upon stimulation with 5nM Wnt3a was unaffected by Clathrin HC knockdown in HEK293T cells. (C) FACS analysis of mESCs stained with OMP-18R5 and detected with Alexa-488 secondary showed that Clathrin HC knockdown does not alter cell surface level of Frizzled receptors. (D) Immunoprecipitation of biotinylated cell surface proteins was followed by quantitative western blot to detect surface Lrp6 with or without 30-minute stimulation with 5nM Wnt3a. The cytoplasmic control GAPDH was detected in lysates but not in immunoprecipitated cell surface fractions. Bar graph on the right shows surface Lrp6 signal normalized by total Lrp6 signal. Clathrin HC knockdown did not lead to a significant change in surface Lrp6 level following 30-minute Wnt3a stimulation. (E) Immunoprecipitation of biotinylated cell surface proteins was followed by quantitative western blot to detect surface Lrp6 with or without 8-hour stimulation with 5nM Wnt3a. Clathrin HC knockdown did not lead to a significant change in surface Lrp6 level following 8-hour Wnt3a stimulation.

### Clathrin-mediated Endocytosis Is Not a Significant Modulator of Wnt Receptor Abundance on the Plasma Membrane

It has been suggested that endocytosis plays a role in downregulation of Wnt signaling, either following Wnt ligand-receptor interaction or without it (15). Endocytic internalization of Wnt receptors such as Frizzled and Lrp5/6 could dampen Wnt activity by reducing the amount of receptors available for Wnt binding on the plasma membrane. To determine whether Clathrin-mediated endocytosis down-regulates Wnt signaling through receptor internalization, we quantified membrane Frizzled abundance via FACS analysis and membrane Lrp6 abundance via surface biotinylation in cells with impaired Clathrin endocytosis.

To quantify Frizzled abundance, we used OMP-18R5, a monoclonal antibody engineered to bind Frizzled 1, 2, 5, 7, and 8 at their extracellular Wnt-interacting domain (34). OMP-18R5 allowed us to concurrently assess multiple Frizzled receptors that are among the most highly expressed in mESCs (32). If Clathrin-mediated endocytosis functions as a constitutive modulator of membrane Frizzled, cells with impaired Clathrin endocytosis would display higher Frizzled signal intensity. However, FACS analysis showed that knock-down of Clathrin HC does not affect Frizzled level on the cell surface (Fig 4C).

For quantification of membrane Lrp6, cell surface proteins were biotinylated and immunoprecipitated. Lrp6 level in the immunoprecipitated surface fraction was measured in a quantitative Western blot, and the surface signal was normalized by whole cell Lrp6 signal of each sample. Normalized levels of Lrp6 demonstrated no significant change in surface presentation of Lrp6 in cells with reduced Clathrin HC (Fig 4D). Membrane Lrp6 level was also surveyed following a 30-minute or 8-hour treatment with Wnt3a to determine whether Wnt stimulation induces internalization of Lrp6. Quantitative Western blot in these contexts showed no notable difference in membrane Lrp6 level between control cells and cells with impaired Clathrin-mediated endocytosis (Fig 4D and 4E). Along with the Frizzled FACS experiments, these results demonstrated that Clathrin-mediated endocytosis does not significantly contribute to downregulation of Wnt receptors on the plasma membrane.

### Monodansylcadaverine Blocks Wnt Signaling Independent of Endocytosis

Some past studies of the role of endocytosis in Wnt signaling relied on the use of a small molecule, monodansylcadaverine (MDC), which reportedly inhibits Clathrin-mediated endocytosis (11, 13, 15). To evaluate the effect of this inhibitor on Wnt signaling, *β*-catenin knock-in reporter mESCs were stimulated with Wnt3a in the presence of MDC. Unlike genetic disruption of Clathrin-mediated endocytosis, incubation with MDC suppressed *β*-catenin accumulation in the nucleus (Fig 5A). Consistent with this, MDC inhibited Wnt3a-induced Axin2 transcription in mESCs and Wnt3a-induced activity of the TOPFlash reporter in mouse L cells in a dose-dependent manner (Fig 5B).

**Fig. 5.**
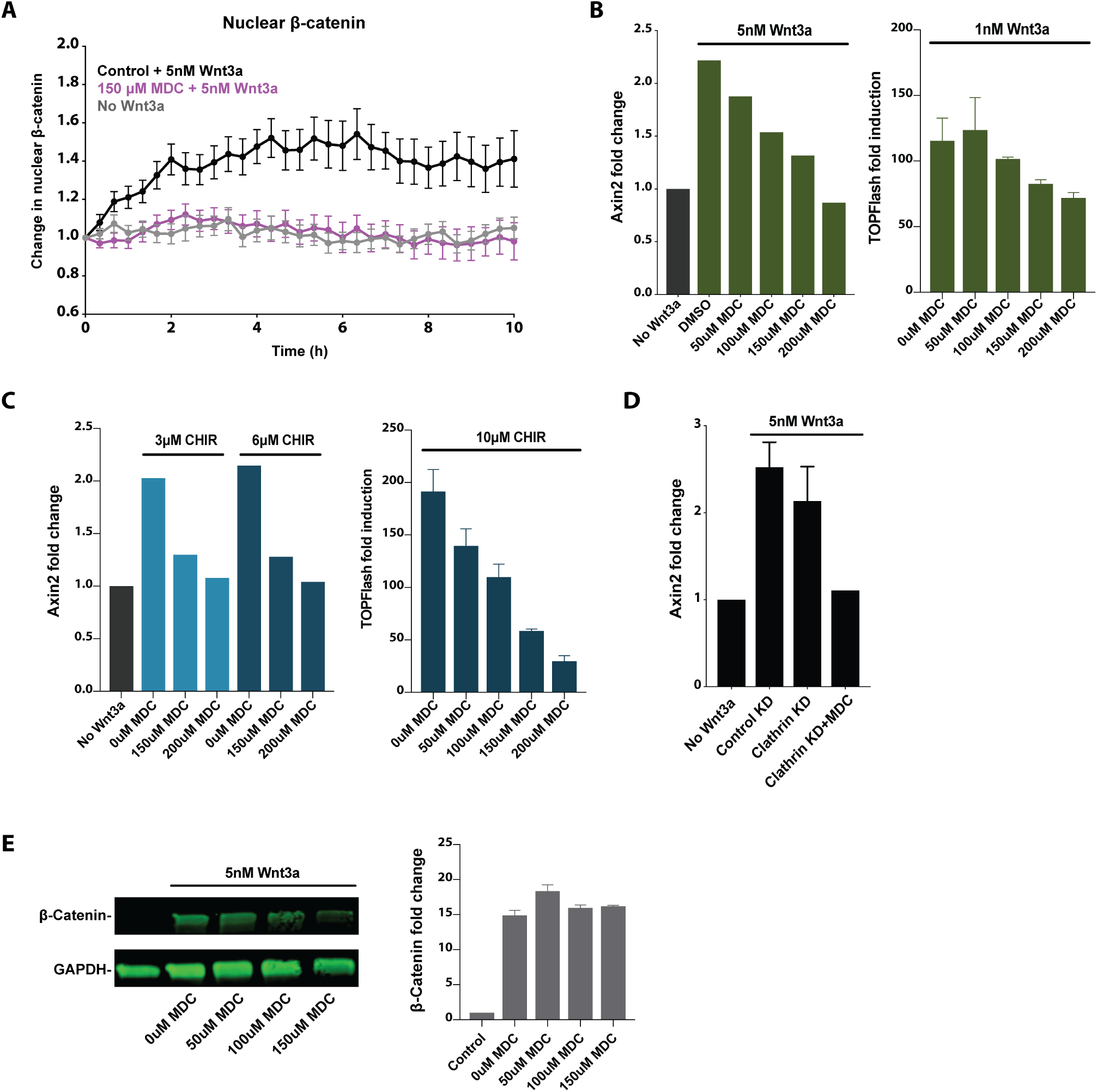
Monodansylcadaverine blocks Wnt signaling independent of endocytosis. (A) Wnt3a induced nuclear translocation of *β*-catenin, which was blocked by addition of 150*μ*M MDC. (Wnt3a, n=33; Wnt3a with 150*μ*M MDC, n=21; no Wnt3a, n=30). (B) MDC impaired Wnt3a-induced transcription of Axin2 in mESCs and Wnt3a-induced TOPFlash luciferase activity in LS/L cells in a dose-dependent manner. (C) MDC impaired CHIR99021-induced transcription of Axin2 in mESCs and CHIR99021-induced TOPFlash luciferase activity in LS/L cells in a dose-dependent manner. (D) MDC blocked Wnt3a-induced Axin2 transcription in mESCs subjected to Clathrin HC knockdown. (E) Western blot showed accumulation of *β*-catenin in L cells stimulated with 5nM Wnt3a for 2 hours, which was not significantly reduced by MDC treatment. Bar graph on the right shows *β*-catenin signal normalized by GAPDH loading control signal, averaged from two biological replicates.

Unexpectedly, we found that Wnt pathway activation by the GSK3*β* inhibitor CHIR99021 was also blocked by MDC. MDC inhibited Axin2 transcription in mESCs or TOPFlash induction in mouse L cells stimulated with CHIR99021 in a dose-dependent manner (Fig 5C). This indicated that MDC affects Wnt signal transduction downstream of GSK3*β* inhibition, not as an endocytosis inhibitor as previously reported. In agreement with this, MDC blocked Wnt-induced Axin2 transcription in mESCs in which Clathrin-mediated endocytosis was already impaired (Fig 5D). Accumulation of cellular *β*-catenin was not blocked by MDC, as measured by Western blot of mouse L cells following stimulation with Wnt3a (Fig 5E). Altogether, these results support that inhibition of Wnt signal transduction by MDC, a small molecule commonly used to block Clathrin-mediated endocytosis, is likely due to an independent, unknown effect on the pathway downstream of GSK3 inhibition and cellular *β*-catenin accumulation. This unexpected effect of MDC might be one source of the discrepancy among reports on the role of endocytosis in Wnt signaling.

## Discussion

Conflicting reports in the field and potential involvement of receptor endocytosis as an essential step in signal transduction warranted a careful examination of the role of endocytosis in Wnt signaling. In order to determine the role of Clathrin-mediated receptor endocytosis in Wnt signal transduction, we used quantitative cell biological and biochemical tools to evaluate the effect of genetic disruption of Clathrin-mediated endocytosis on downstream events in Wnt signal transduction.

Our results led to the conclusion that endocytosis is not a general requirement for Wnt signal transduction. First, we confirmed that a mESC line reports pathway activation at the level of *β*-catenin accumulation in the cytoplasm and the nucleus in a Wnt-specific, temporally sensitive manner. Knockout or knockdown of essential components of Clathrin-mediated endocytosis such as Clathrin HC or AP2 subunits in these mESCs severely impaired their uptake of the Clathrin cargo Transferrin. However, Wnt signal transduction was unaffected by these disruptions, as evinced by appropriate increase in cytoplasmic and nuclear *β*-catenin and target gene induction. Furthermore, cells with impaired Clathrin function maintained their pluripotency, a key biological consequence of Wnt pathway activation in mESCs. Lack of Wnt signaling phenotype in additional mouse and human cell lines provided further support that Clathrin-mediated endocytosis is not required for Wnt pathway activation. Suppression of Caveolin-mediated endocytosis, which has also been implicated in Wnt receptor endocytosis, did not compromise Wnt signal transduction in mESCs.

These results provide ample evidence that receptor endocytosis is not an essential event in Wnt pathway activation. In addition, they challenge an existing Wnt signal transduction model that involves MVBs (19). In this model, endocytic structures containing Wnt pathway proteins including GSK3*β* mature into MVBs with intralumenal vesicles. MVBs sequester GSK3*β* away from cytoplasmic *β*-catenin, leading to *β*-catenin stabilization and accumulation. However, this model requires receptor endocytosis for subsequent MVB formation and Wnt pathway activation.

Additional lines of evidence in the field support that internalization of the ligand and the receptor complex via endocytosis is not a general requirement for Wnt pathway activation. There are instances in which overexpression or manipulation of Wnt pathway components within the cytoplasm is sufficient to initiate signaling: overexpression of the cytoplasmic Wnt factor Dishevelled (Dvl) or optogenetic oligomerization of the cytoplasmic portion of Lrp6 (35) (36). Second, exposure to Wnt molecules covalently attached to a magnetic bead that therefore cannot be internalized by cells is sufficient for pathway activation in mESCs (37). Third, super-resolution microscopy of GFP-tagged Dvl2 revealed its undirected movement and lack of colocalization with endocytic vesicle markers. These results challenge models in which pathway activation involves internalization of receptor complexes associated with Dvl (38).

There are several potential sources of discrepancy between results of this study and some previous reports in the field. Among these are pleiotropic effects of disrupting endocytic factors. One example is Clathrin’s role in mitosis, which is independent of its function in endocytosis. Consistent with Clathrin’s reported role in mitotic spindle organization (29), knockdown of Clathrin HC led to reduced cell proliferation. In addition, Gagliardi and colleagues reported a decrease in levels of multiple Wnt pathway proteins with or without Wnt stimulation in the presence of endocytosis inhibitors and suggested that inhibition of endocytosis may trigger degradation of multiple proteins through an unknown mechanism (39). Finally, disruption of Dynamin, which has been employed to inhibit endocytosis, could lead to pleiotropic effects and confound study conclusions. Dynamin is responsible for pinching off vesicles from the Golgi network as well as from the plasma membrane (40) (41). Secretory pathways as well as Clathrin- and Caveolin-mediated endocytic pathways could be affected in cells with impaired Dynamin function. This would make it difficult to separate the effect of Dynamin disruption on Wnt ligand secretion from that on ligand uptake, both of which could attenuate Wnt signaling.

To address pleiotropic consequences of endocytosis disruption, it is important to carefully control for the quantity and quality of samples with or without endocytosis suppression when assaying for Wnt pathway activation. In addition, endocytic genes should be disrupted specifically in Wnt-receiving cells. To this end, we utilized single mESCs removed from other cell types, which allowed us to probe the effects of endocytosis disruption on single Wnt receiving cells.

In addition, the use of overexpressed or non-native Wnt pathway components might have led to disagreements among study conclusions. In some studies, a truncated, constitutively active form of Lrp6 was expressed in order to activate Wnt signaling rather than adding a purified Wnt ligand or media containing Wnt ligand (19, 42). In others, assessment of receptor internalization and Wnt signal transduction was conducted with overexpressed, tagged receptors (11, 43). As overexpressed or non-native Wnt pathway proteins might be endocytosed or trafficked differently, endogenous Wnt path-way components are better suited for investigating the mechanism of signal transduction. Therefore, we relied on purified Wnt ligand for pathway activation and accumulation of endogenous *β*-catenin and target gene transcription for signal transduction readouts.

A third possible source of discrepancy is non-specific effects of endocytosis inhibitors. Some previous efforts relied heavily on the small molecule inhibitor MDC to suppress Clathrin-mediated endocytosis (11, 13, 15, 44). Our evaluation at different points along Wnt signal transduction showed that MDC affects the Wnt pathway downstream of GSK3*β* inhibition. MDC inhibited Wnt-induced target transcription in mESCs in which Clathrin-mediated endocytosis was already impaired, further supporting that MDC suppresses Wnt signaling independent of endocytosis. Future endeavors to study the mechanism of Wnt signal transduction should be wary of off-target actions of small molecule inhibitors as well as pleiotropic effects of disrupted genes and behavior of over-expressed proteins.

It should be noted that Wnt signal transduction could involve receptor endocytosis under specific circumstances. A recent study identified one such context in which EGFR functions as a co-receptor to facilitate Wnt9a-Fzd9b specific interaction in hematopoiesis (45). In this context, Clathrin-mediated endocytosis and subsequent activation of *β*-catenin-dependent Wnt signaling were required for differentiation of zebrafish and human hematopoietic stem and progenitor cells. This is consistent with other accounts of endocytosis induction upon EGFR activation, and leaves open the possibility that Wnt pathway activation facilitated by unconventional co-receptors depends on receptor endocytosis.

We also addressed involvement of Clathrin-mediated endocytosis as a negative regulator of Wnt signaling through receptor downregulation. Clathrin HC knockdown or AP2*α* knockout did not alter cell surface levels of Frizzled and Lrp6. Even with Wnt3a stimulation, disruption of Clathrin endocytosis did not lead to a detectable increase in surface Lrp6 level. This is consistent with a previous observation that Wnt stimulation does not induce Lrp6 internalization (14). Together, results from these experiments indicate that Clathrin-mediated endocytosis does not play a significant role in constitutive or Wnt-induced receptor downregulation.

In summary, requirement for endocytosis in Wnt signal transduction was tested in an investigation that combined a Wnt-responsive *in vitro* system, genetic approaches to target essential components of endocytosis, and means to monitor endogenous Wnt signal transducers in single cells to assess pathway activation. We conclude that receptor endocytosis is not required for Wnt signal transduction.

## Materials and Methods

### Cell culture and reagents

R1 mESCs were purchased from ATCC and cultured at 37°C and 5% CO_2_. Routine culture of mESCs was carried out in medium containing DMEM with GlutaMAX (Thermo Fisher, Waltham, MA), 15% fetal bovine serum (GE Healthcare, Chicago, IL), 1 mM sodium pyruvate, MEM non-essential amino acids, 50 *μ*M *β*-mercaptoethanol, 100 U/ml penicillin, 100 *μ*g /ml streptomycin (all from Thermo Fisher), and 1,000 U/ml LIF (Millipore Sigma, Burlington, MA) in tissue culture plates coated with 0.1% gelatin. For serum-free culture, cells were maintained in one volume DMEM/F12 with GlutaMAX combined with one volume Neurobasal medium, with 0.5% N2 Supplement, 1% B27 Supplement, 0.033% BSA, 50 *μ*M *β*-mercaptoethanol, 100 U/ml penicillin and 100 *μ*g/ml streptomycin (all from Thermo Fisher) in tissue culture plates coated with 2.5*μ*g/cm^2^ human fibronectin (Thermo Fisher). HEK293T cells and L cells were cultured in DMEM with GlutaMAX supplemented with 10% fetal bovine serum (Omega Scientific, Tarzana, CA), 100 U/ml penicillin, and 100 *μ*g/ml streptomycin. Career-free recombinant mouse Wnt3a was purchased from R&D, reconstituted in 0.1% BSA/PBS, and added to culture at specified concentrations.

### DNA constructs

mESC line harboring a *β*-catenin knock-in reporter, with the C-terminus of *β*-catenin tagged with the yellow fluorescent protein Venus, was a kind gift from Alexander Aulehla at EMBL Heidelberg. In order to add a nucleus marker to the reporter mESCs, CSII-EF1-MCS vector encoding the histone protein H2B tagged with the cyan fluorescent protein mTurquoise was introduced via Lentiviral infection.

### CRISPR-mediated knockout

For CRISPR-mediated knockout of AP2*α*, sgRNAs targeting exon 4 of mouse AP2a1 were introduced into a vector encoding Cas9 from *S. pyogenes* with 2A-Puro selection cassette, pSpCas9(BB)-2A-Puro v2.0 (Plasmid #62988 from Addgene). sgRNAs were designed using CHOPCHOP v2 (46). sgRNA-mediated cutting efficiency at target genomic site was evaluated with Surveyor Mutation Detection Kit (Integrated DNA Technologies, Coralville, IA). mESC transfection with pSpCas9(BB)-2A-Puro v2.0 containing confirmed sgRNAs was performed using Effectene Transfection Reagent (Qiagen, Germantown, MD). After clonal selection, colonies were tested for loss of AP2*α* through genotyping and Western blotting. Genotyping analysis was assisted with the web tool TIDE: Tracking of Indels by DEcomposition (47).

### Western blots

Cells were lysed in RIPA buffer [150 mM NaCl, 1% IGEPAL® CA-630, 0.5% of Sodium Deoxycholate, 0.1% sodium dodecylsulfate (SDS), and 50mM Tris] supplemented with protease inhibitor cocktail (Biovision, Milpitas, CA). Total protein concentration of the lysates was measured using Bicinchoninic Acid (BCA) Assay (Thermo Fisher) with bovine serum albumin as a standard. Proteins were resolved via SDS-PAGE in 10% Mini-PROTEAN Tris-Glycine gels (Bio-Rad, Hercules, CA) and wet transferred to 0.45 *μ*m nitrocellulose membrane (GE Healthcare). Membranes were blocked with either 5% milk dissolved in TBS (Tris buffered saline) or Intercept TBS blocking buffer (LI-COR Biosciences, Lincoln, NE) for 1 hour at room temperature then incubated overnight at 4°C with primary antibodies diluted in the blocking solution containing 0.1% Tween-20. Membranes were washed in TBST and incubated with HRP- or IRDye-conjugated secondary antibody diluted at 1:5000 in blocking solution containing 0.1% Tween-20. After washing, protein signals were detected via enhanced chemiluminescence (Western Lightning Plus ECL, Perkin Elmer, Waltham, MA) followed by exposure to film (CL-XPosure, Thermo Fisher). For quantitative blots, protein signals were detected on LI-COR Odyssey CLx and quantified using Empiria Studio software.

### Alkaline phosphatase stain

mESCs were cultured in serum-free mESC media with or without Wnt3a or CHIR99021 for 4 days at low to medium density. Alkaline phosphatase activity was detected using the Alkaline Phosphatase Detection Kit (Millipore Sigma) according to the manufacturer’s protocol. mESC colonies were fixed with 4% paraformaldehyde in PBS for 2 min and rinsed with TBST (Tris buffered saline, 0.1% Tween-20). Cells were then stained with alkaline phosphatase solution at room temperature for 15 min in the dark and imaged in PBS after another TBST wash. Mean stain intensity for each mESC colony was quantified using ImageJ following background subtraction.

### siRNAs and antibodies

Following antibodies were used for Western blots: *β*-catenin (BD Biosciences, San Jose, CA; BD 610154; 1:1000), Lrp6 (Cell Signaling Technology, Danvers, MA; CST 3395; 1:1000), Clathrin heavy chain (Santa Cruz Biotechnology, Dallas, TX; SC-12734; 1:1000), AP-2 subunit *α* (BD 610501; 1:1000), AP-2 subunit *μ*2 (BD 611350; 1:1000), *β*-tubulin (SC-9104; 1:1000), GAPDH (Proteintech, Rosemont, IL; 60004-1; 1:2000). Following siRNAs from Dharmacon (GE Healthcare) were used for siRNA-mediated knock-down: siGenome Mouse Lrp6 (16974) siRNA SMART-pool, siGenome Mouse Cltc (67300) siRNA SMART-pool, siGenome Mouse Cav1 (12389) siRNA SMART-pool, siGenome Mouse Cav2 (12390) siRNA SMART-pool, siGenome Mouse Ap2m1 (11773) siRNA SMARTpool, siGenome Human Cltc (1213) siRNA SMARTpool, Custom control siRNA (5’-CGCACGUAUAUGACAAUCGUU-3’).

### Live cell microscopy and analysis

All time-lapse microscopy experiments were performed using Zeiss Cell Observer Spinning Disk confocal system. *β*-catenin reporter mESCs were adjusted to a Wnt-inactive state for 24 h in media without 2i or Wnt3a. Single cell suspension of these reporter mESCs were plated in Nunc 8-well chambered cover-glass slides (Thermo Fisher) coated with 2.5*μ*g/cm^2^ human fibronectin at 10^4^ cells per well density. Cells were allowed to adhere for 2 h at 37°C and 5% CO_2_. During time-lapse experiments, DIC and fluorescence images were acquired every 20 min using a 20x/NA 0.8 air objective, Evolve 512 EMCCD camera, and Zeiss Zen Blue software. Individual time frame images in each channel were exported for subsequent analysis in Matlab 9.2 (Mathwork). Cells undergoing division or in contact with other cells were excluded from analysis. For each time frame, a cell mask was created using the DIC image and a nucleus mask was created using the 405nm channel image. Cellular and nuclear signal intensity of *β*-catenin, measured in the 488nm channel, was quantified using these cell and nucleus masks, respectively. Change in cellular *β*-catenin signal and change in nuclear *β*-catenin signal relative to the first time point were tracked for each cell. Prism 8 (GraphPad) was used to plot the average and standard deviation of these relative changes for all cells, and to perform Student’s-t test to examine the difference between control and endocytosis perturbed conditions.

### Luciferase reporter assay

*β*-catenin mediated transcription activation was determined using mouse L cells stably expressing SuperTOPFlash (firefly luciferase reporter driven by 7x TCF/LEF *β*-catenin binding sites) and *β*-galactosidase for normalization. Cells were seeded at 5×10^4^ cells per well in 96-well culture plates, incubated overnight with Wnt3a or CHIR99021 at specified concentrations, then lysed in passive lysis buffer (Promega, Madison, WI) for reporter assay. Relative luciferase activity units were measured and normalized against *β*-galactosidase activity using the Dual-Light assay system (Thermo Fisher) on LB 960 Centro luminometer (Berthold Technologies, Oak Ridge, TN). All assays were performed in triplicates.

### Reverse transcription and quantitative real-time PCR

To activate Wnt signaling, cells were stimulated for specified amount of time with 1nM-5nM Wnt3a in medium. Total RNA was prepared using RNeasy mini kit (Qiagen) according to the manufacturer’s protocol. RNA was reverse transcribed using random primers (High Capacity cDNA Reverse Transcription kit, Applied Biosystems, Foster City, CA). Gene expression was assayed by real-time PCR using TaqMan Gene Expression Assays (Applied Biosystems) on Applied Biosystems StepOnePlus real-time PCR system. All PCRs were carried out in triplicates with glyceraldehyde-3-phosphate dehydrogenase (GAPDH) as a reference gene. The following TaqMan probes were used: mouse Axin2 (Mm00443610), mouse Caveolin1 (Mm00483057), mouse Caveolin2 (Mm01129337), mouse Clathrin heavy chain (Mm01307036), mouse GAPDH (Mm99999915), human Axin2 (Hs00610344), and human GAPDH (4326317E).

### Transferrin uptake assay

Prior to assay, cells were cultured in serum-free media on coverslips for 24 h. Cells were pulsed with 25ug/mL Transferrin-488 (Thermo Fisher T13342) in serum-free media on ice for 30 min. Coverslips with cells were transferred to 37°C for 20 min induce Transferrin uptake while control coverslips were kept at 4°C. After PBS washes, cells were rinsed for 40 sec with acidic stripping solution [500mM NaCl and 100mM glacial acetic acid in distilled water] to remove surface-bound Transferrin. This was followed by PBS washes and fixing with 3.2% paraformaldehyde in serum-free media at 4°C for 20 min. After PBS washes, coverslips were mounted in Prolong Gold Antifade reagent with DAPI (Thermo Fisher P36930), dried overnight, and imaged.

### Fluorescence-activated cell sorting

mESCs were harvested with Accutase (Thermo Fisher) and washed with FACS buffer (2% FBS in PBS) at 4°C. Primary antibody staining was performed on ice for 1 hour followed by a FACS buffer wash and secondary antibody incubation on ice for 45 min in dark. To exclude dead cells, samples were incubated with 5*μ*g/mL DAPI (Thermo Fisher D1306) in FACS buffer on ice for 15 min. Cells were washed, resuspended in FACS buffer, and analyzed on LSRFortessa (BD Biosciences) or sorted on FACSAria IIu (BD Biosciences). Data were processed with FACSDiva software and FlowJo v10. Doublets were excluded by FSC-W × FSC-H and SSC-W × SSC-H analysis, and dead cells were excluded by DAPI detection. The following antibodies were used: OMP-18R5 (OncoMed Pharmaceuticals, Redwood City, CA) at 10*μ*g/mL with antihuman IgG (Fc) secondary conjugated to FITC (Millipore Sigma AP113F).

### Cell surface protein biotinylation

For quantification of surface Lrp6, cell surface proteins were biotinylated using EZ-Link Sulfo-NHS-SS-Biotin (Thermo Fisher) according to the manufacturer’s protocol. Surface proteins were labeled with 800*μ*M Sulfo-NHS-SS-Biotin in PBS for 30 min at 4°C. Cells were washed twice with ice-cold 20mM Tris buffer to quench non-reacted biotinylation reagent then with ice-cold PBS. Cells were lysed in RIPA buffer supplemented with protease inhibitor cocktail (Biovision) and 1mM PMSF. Total protein concentration of the lysates was measured via BCA assay and biotinylated proteins were immunoprecipitated using Streptavidin magnetic beads (Thermo Fisher). 500*μ*g magnetic beads were incubated with up to 70*μ*g of lysate rotating overnight at 4°C, washed with RIPA buffer, and eluted in 1x Laemmli sample buffer supplemented with 2mM Biotin and 20mM DTT by boiling for 3 min at 95°C. Quantitative Western blot was performed with elutes and whole cell lysates as described above and imaged on LI-COR Odyssey CLx. Lrp6 antibody was a kind gift from Gary Davidson.

## ACKNOWLEDGEMENTS

We thank Alexander Aulehla and Jana Kress for generating and sharing *β*-catenin knock-in reporter mESCs. We thank Catriona Logan and Teni Anbarchian for feedback on the study and the manuscript, and Wenzhe Ma and Alexander Aulehla for fruitful discussions. We acknowledge Stanford Neuroscience Microscopy Service, supported by NIH NS069375, for assistance with initial imaging data collection. EYR was supported by a Stanford Graduate Fellowship.

## Supplementary Figures

**Fig. 6.**
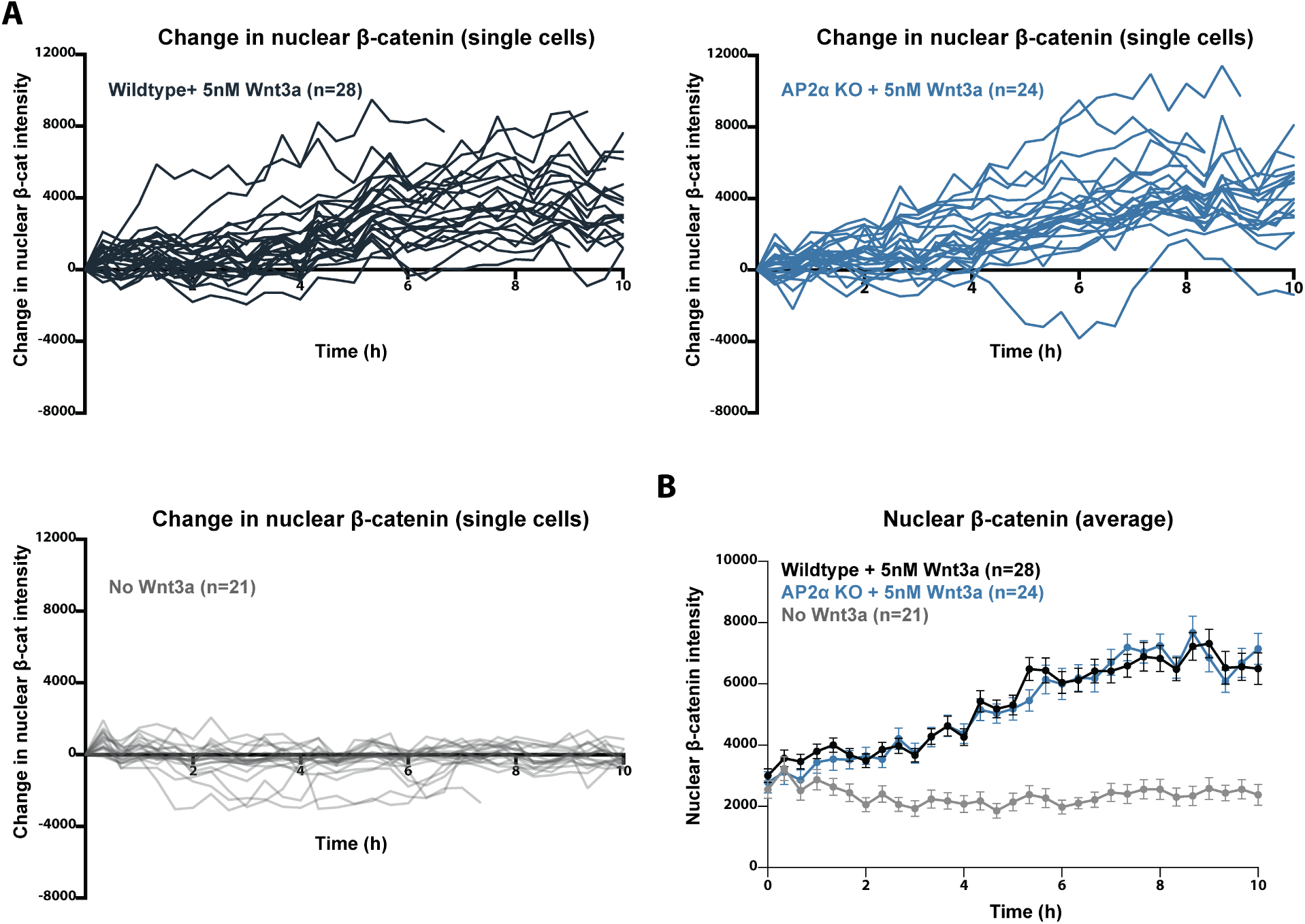
Loss of AP2*α* does not affect absolute level changes in nuclear *β*-catenin. (A) Change in absolute levels of nuclear *β*-catenin was quantified in wildtype cells and AP2*α* knockout cells with 5nM Wnt3a. Each track represents a cell imaged over 10 hours. (Wildtype, n=28; AP2*α* knockout, n=24; no Wnt3a, n=21). (B) Change in absolute levels of nuclear *β*-catenin shown in (A) was averaged.

**Fig. 7.**
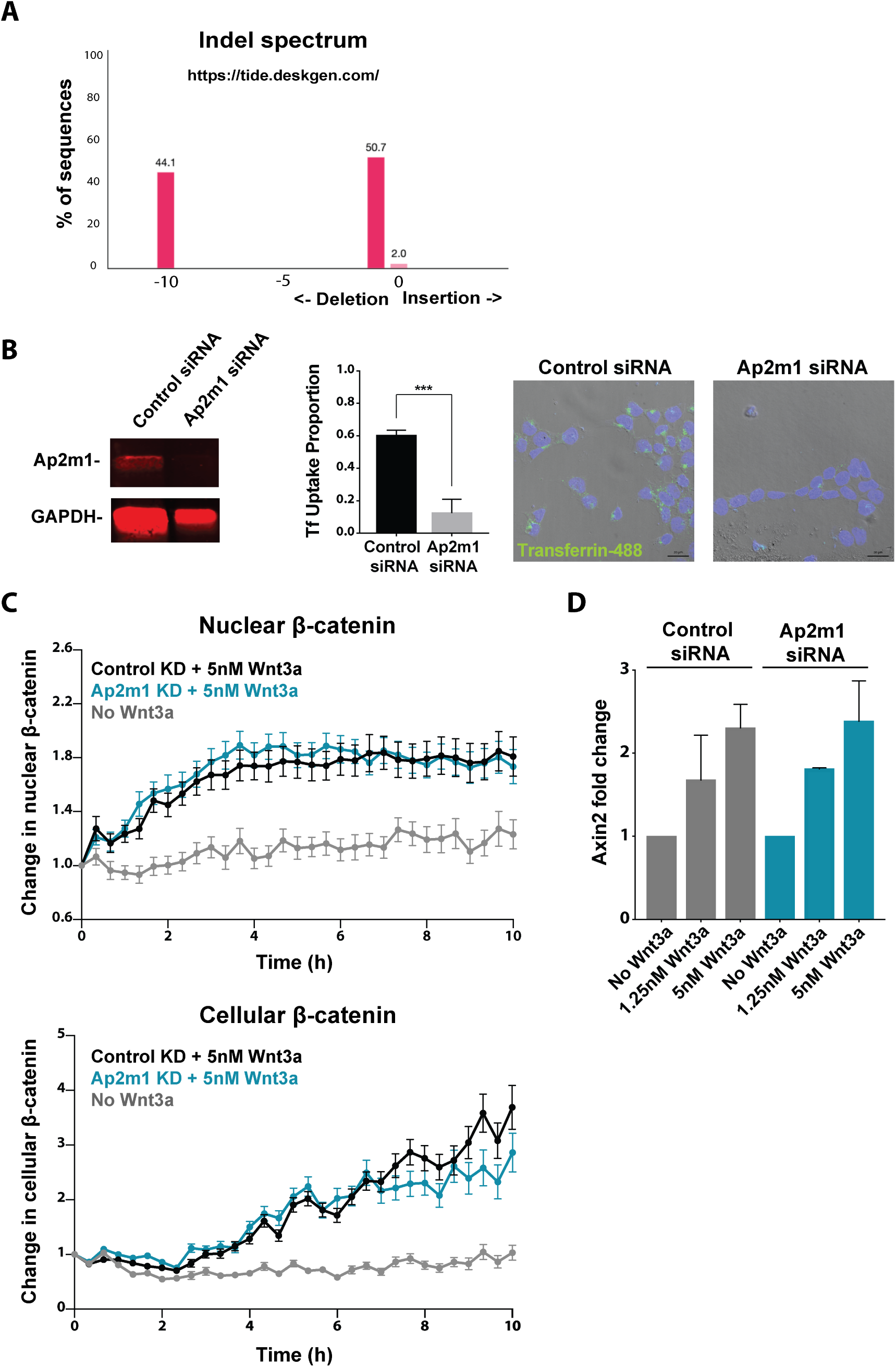
Knockdown of AP2*μ* impairs Clathrin-mediated cargo uptake but does not affect Wnt signal transduction. (A) Following CRISPR-mediated targeting of AP2*α*, a clone with frameshift deletion alleles was identified. (B) Effective siRNA-mediated knockdown of AP2*μ* was confirmed via Western blot. AP2*μ* knockdown impaired cellular uptake of fluorescently labeled Transferrin (p *<* 0.001). (C) AP2*μ* knockdown did not affect nuclear translocation or cellular accumulation of *β*-catenin upon stimulation with 5nM Wnt3a. (Control siRNA, n=30; AP2*μ* siRNA, n=26; no Wnt3a, n=17). (D) Transcription of the Wnt target gene Axin2 was unaffected by knockdown of AP2*μ*.

**Fig. 8.**
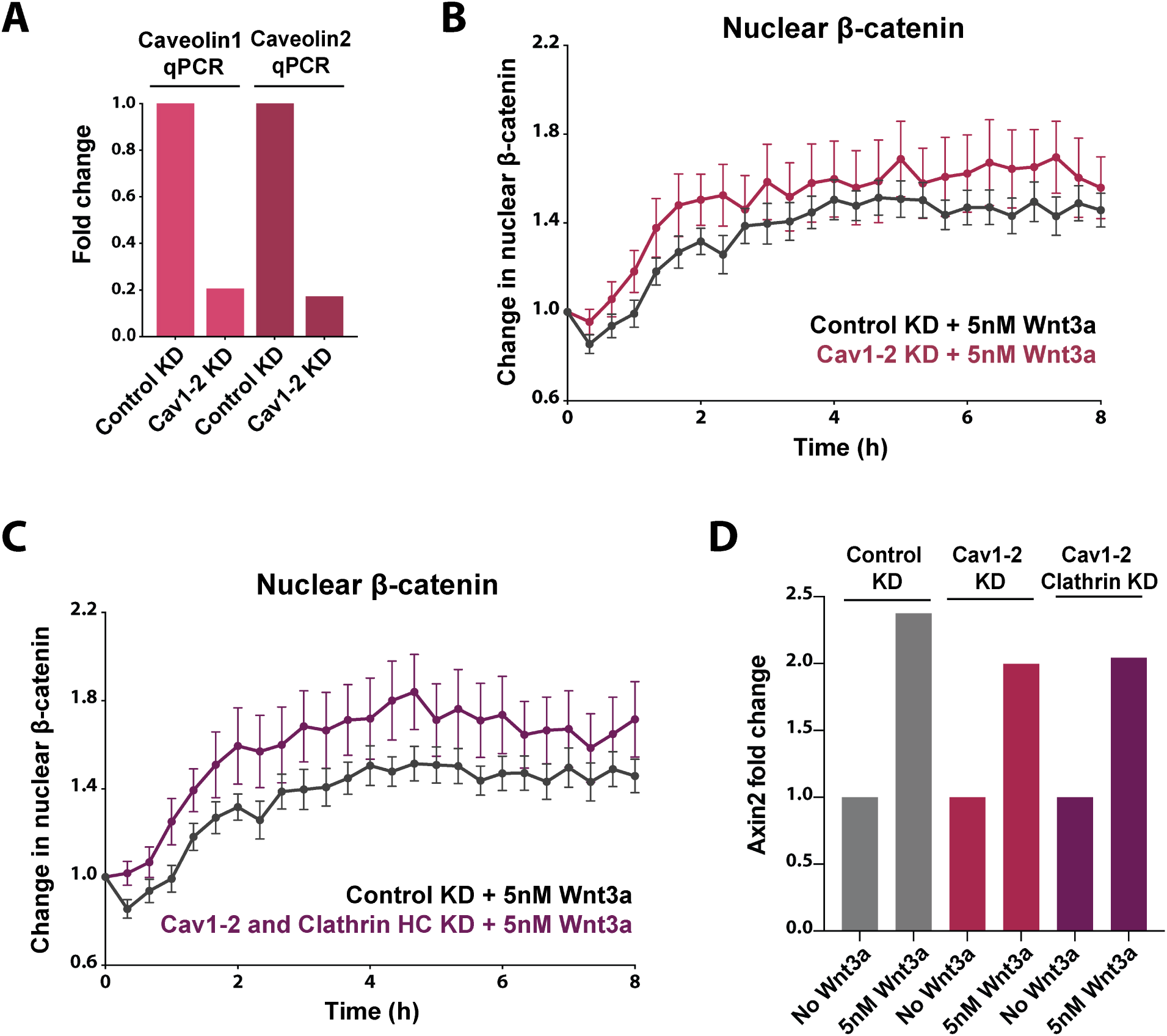
Double knockdown of Caveolin1 and 2 does not affect Wnt signal transduction. (A) Simultaneous knockdown of Caveolin1 and Caveolin2 was confirmed via qPCR. (B) Caveolin 1 and 2 double knockdown did not affect nuclear translocation of *β*-catenin upon stimulation with 5nM Wnt3a (Control siRNA, n=19; Caveolin1-2 siRNA, n=14). (C) Triple knockdown of Caveolin 1-2 and Clathrin HC did not affect nuclear translocation of *β*-catenin upon stimulation with 5nM Wnt3a (Control siRNA, n=19; Caveolin1-2 and Clathrin HC siRNA, n=14). (D) Transcription of the Wnt target gene Axin2 was unaffected by double knockdown of Caveolin 1 and 2 or triple knockdown of Caveolin 1-2 and Clathrin HC.

## Bibliography

1. K. M. Loh, R. van Amerongen, and R. Nusse. Generating cellular diversity and spatial form: Wnt signaling and the evolution of multicellular animals. Dev Cell, 38(6):643–55, 2016. ISSN 1878-1551 (Electronic) 1534-5807 (Linking). doi: 10.1016/j.devcel.2016.08.011.

2. S. Rudloff and R. Kemler. Differential requirements for beta-catenin during mouse development. Development, 139(20):3711–21, 2012. ISSN 1477-9129 (Electronic) 0950-1991 (Linking). doi: 10.1242/dev.085597.

3. C. Y. Logan and R. Nusse. The wnt signaling pathway in development and disease. Annu Rev Cell Dev Biol, 20:781–810, 2004. ISSN 1081-0706 (Print) 1081-0706 (Linking). doi: 10.1146/annurev.cellbio.20.010403.113126.

4. K. Willert, J. D. Brown, E. Danenberg, A. W. Duncan, I. L. Weissman, T. Reya, 3rd Yates, J. R., and R. Nusse. Wnt proteins are lipid-modified and can act as stem cell growth factors. Nature, 423(6938):448–52, 2003. ISSN 0028-0836 (Print) 0028-0836 (Linking). doi: 10.1038/nature01611.

5. A. Ring, Y. M. Kim, and M. Kahn. Wnt/catenin signaling in adult stem cell physiology and disease. Stem Cell Rev Rep, 10(4):512–25, 2014. ISSN 2629-3277 (Electronic). doi: 10.1007/s12015-014-9515-2.

6. T. Reya and H. Clevers. Wnt signalling in stem cells and cancer. Nature, 434(7035):843–50, 2005. ISSN 1476-4687 (Electronic) 0028-0836 (Linking). doi: 10.1038/nature03319.

7. S. H. Tan and N. Barker. Wnt signaling in adult epithelial stem cells and cancer. Prog Mol Biol Transl Sci, 153:21–79, 2018. ISSN 1878-0814 (Electronic) 1877-1173 (Linking). doi: 10.1016/bs.pmbts.2017.11.017.

8. A. R. Hernandez, A. M. Klein, and M. W. Kirschner. Kinetic responses of beta-catenin specify the sites of wnt control. Science, 338(6112):1337–40, 2012. ISSN 1095-9203 (Electronic) 0036-8075 (Linking). doi: 10.1126/science.1228734.

9. M. van Noort, J. Meeldijk, R. van der Zee, O. Destree, and H. Clevers. Wnt signaling controls the phosphorylation status of beta-catenin. J Biol Chem, 277(20):17901–5, 2002. ISSN 0021-9258 (Print) 0021-9258 (Linking). doi: 10.1074/jbc.M111635200.

10. A. Tomas, C. E. Futter, and E. R. Eden. Egf receptor trafficking: consequences for signaling and cancer. Trends Cell Biol, 24(1):26–34, 2014. ISSN 1879-3088 (Electronic) 0962-8924 (Linking). doi: 10.1016/j.tcb.2013.11.002.

11. H. Yamamoto, H. Komekado, and A. Kikuchi. Caveolin is necessary for wnt-3a-dependent internalization of lrp6 and accumulation of beta-catenin. Dev Cell, 11(2):213–23, 2006. ISSN 1534-5807 (Print) 1534-5807 (Linking). doi: 10.1016/j.devcel.2006.07.003.

12. E. S. Seto and H. J. Bellen. Internalization is required for proper wingless signaling in drosophila melanogaster. J Cell Biol, 173(1):95–106, 2006. ISSN 0021-9525 (Print) 0021-9525 (Linking). doi: 10.1083/jcb.200510123.

13. J. T. Blitzer and R. Nusse. A critical role for endocytosis in wnt signaling. BMC Cell Biol, 7: 28, 2006. ISSN 1471-2121 (Electronic) 1471-2121 (Linking). doi: 10.1186/1471-2121-7-28.

14. I. Kim, W. Pan, S. A. Jones, Y. Zhang, X. Zhuang, and D. Wu. Clathrin and ap2 are required for ptdins(4,5)p2-mediated formation of lrp6 signalosomes. J Cell Biol, 200(4):419–28, 2013. ISSN 1540-8140 (Electronic) 0021-9525 (Linking). doi: 10.1083/jcb.201206096.

15. C. C. Liu, T. Kanekiyo, B. Roth, and G. Bu. Tyrosine-based signal mediates lrp6 receptor endocytosis and desensitization of wnt/beta-catenin pathway signaling. J Biol Chem, 289 (40):27562–70, 2014. ISSN 1083-351X (Electronic) 0021-9258 (Linking). doi: 10.1074/jbc.M113.533927.

16. M. J. Agajanian, M. P. Walker, A. D. Axtman, R. R. Ruela-de Sousa, D. S. Serafin, A. D. Rabinowitz, D. M. Graham, M. B. Ryan, T. Tamir, Y. Nakamichi, M. V. Gammons, J. M. Bennett, R. M. Counago, D. H. Drewry, J. M. Elkins, C. Gileadi, O. Gileadi, P. H. Godoi, N. Kapadia, S. Muller, A. S. Santiago, F. J. Sorrell, C. I. Wells, O. Fedorov, T. M. Willson, W. J. Zuercher, and M. B. Major. Wnt activates the aak1 kinase to promote clathrin-mediated endocytosis of lrp6 and establish a negative feedback loop. Cell Rep, 26(1):79–93 e8, 2019. ISSN 2211-1247 (Electronic). doi: 10.1016/j.celrep.2018.12.023.

17. M. Kaksonen and A. Roux. Mechanisms of clathrin-mediated endocytosis. Nat Rev Mol Cell Biol, 19(5):313–326, 2018. ISSN 1471-0080 (Electronic) 1471-0072 (Linking). doi: 10.1038/nrm.2017.132.

18. S. H. Hong, C. L. Cortesio, and D. G. Drubin. Machine-learning-based analysis in genome-edited cells reveals the efficiency of clathrin-mediated endocytosis. Cell Rep, 12(12):2121–30, 2015. ISSN 2211-1247 (Electronic). doi: 10.1016/j.celrep.2015.08.048.

19. V. F. Taelman, R. Dobrowolski, J. L. Plouhinec, L. C. Fuentealba, P. P. Vorwald, I. Gumper, D. D. Sabatini, and E. M. De Robertis. Wnt signaling requires sequestration of glycogen synthase kinase 3 inside multivesicular endosomes. Cell, 143(7):1136–48, 2010. ISSN 1097-4172 (Electronic) 0092-8674 (Linking). doi: 10.1016/j.cell.2010.11.034.

20. H. X. Hao, Y. Xie, Y. Zhang, O. Charlat, E. Oster, M. Avello, H. Lei, C. Mickanin, D. Liu, H. Ruffner, X. Mao, Q. Ma, R. Zamponi, T. Bouwmeester, P. M. Finan, M. W. Kirschner, J. A. Porter, F. C. Serluca, and F. Cong. Znrf3 promotes wnt receptor turnover in an rspondin-sensitive manner. Nature, 485(7397):195–200, 2012. ISSN 1476-4687 (Electronic) 0028-0836 (Linking). doi: 10.1038/nature11019.

21. N. Sato, L. Meijer, L. Skaltsounis, P. Greengard, and A. H. Brivanlou. Maintenance of pluripotency in human and mouse embryonic stem cells through activation of wnt signalling by a pharmacological gsk-3-specific inhibitor. Nat Med, 10(1):55–63, 2004. ISSN 1078-8956 (Print) 1078-8956 (Linking). doi: 10.1038/nm979.

22. D. ten Berge, D. Kurek, T. Blauwkamp, W. Koole, A. Maas, E. Eroglu, R. K. Siu, and R. Nusse. Embryonic stem cells require wnt proteins to prevent differentiation to epiblast stem cells. Nat Cell Biol, 13(9):1070–5, 2011. ISSN 1476-4679 (Electronic) 1465-7392 (Linking). doi: 10.1038/ncb2314.

23. K. F. Kelly, D. Y. Ng, G. Jayakumaran, G. A. Wood, H. Koide, and B. W. Doble. beta-catenin enhances oct-4 activity and reinforces pluripotency through a tcf-independent mechanism. Cell Stem Cell, 8(2):214–27, 2011. ISSN 1875-9777 (Electronic) 1875-9777 (Linking). doi: 10.1016/j.stem.2010.12.010.

24. B. J. Merrill. Wnt pathway regulation of embryonic stem cell self-renewal. Cold Spring Harb Perspect Biol, 4(9):a007971, 2012. ISSN 1943-0264 (Electronic) 1943-0264 (Linking). doi: 10.1101/cshperspect.a007971.

25. L. Goentoro and M.W. Kirschner. Evidence that fold-change, and not absolute level, of betacatenin dictates wnt signaling. Mol Cell, 36(5):872–84, 2009. ISSN 1097-4164 (Electronic) 1097-2765 (Linking). doi: 10.1016/j.molcel.2009.11.017.

26. P. Kafri, S. E. Hasenson, I. Kanter, J. Sheinberger, N. Kinor, S. Yunger, and Y. Shav-Tal. Quantifying beta-catenin subcellular dynamics and cyclin d1 mrna transcription during wnt signaling in single living cells. Elife, 5, 2016. ISSN 2050-084X (Electronic) 2050-084X (Linking). doi: 10.7554/eLife.16748.

27. A. P. Turkewitz and S. C. Harrison. Concentration of transferrin receptor in human placental coated vesicles. J Cell Biol, 108(6):2127–35, 1989. ISSN 0021-9525 (Print) 0021-9525 (Linking). doi: 10.1083/jcb.108.6.2127.

28. M. L. Lepire and C. A. Ziomek. Preimplantation mouse embryos express a heat-stable alkaline phosphatase. Biol Reprod, 41(3):464–73, 1989. ISSN 0006-3363 (Print) 0006-3363 (Linking). doi: 10.1095/biolreprod41.3.464.

29. S. J. Royle, N. A. Bright, and L. Lagnado. Clathrin is required for the function of the mitotic spindle. Nature, 434(7037):1152–7, 2005. ISSN 1476-4687 (Electronic) 0028-0836 (Linking). doi: 10.1038/nature03502.

30. S. J. Royle. The role of clathrin in mitotic spindle organisation. J Cell Sci, 125(Pt 1):19–28, 2012. ISSN 1477-9137 (Electronic) 0021-9533 (Linking). doi: 10.1242/jcs.094607.

31. A. G. Wrobel, Z. Kadlecova, J. Kamenicky, J. C. Yang, T. Herrmann, B. T. Kelly, A. J. McCoy, P. R. Evans, S. Martin, S. Muller, F. Sroubek, D. Neuhaus, S. Honing, and D. J. Owen. Temporal ordering in endocytic clathrin-coated vesicle formation via ap2 phosphorylation. Dev Cell, 50(4):494–508 e11, 2019. ISSN 1878-1551 (Electronic) 1534-5807 (Linking). doi: 10.1016/j.devcel.2019.07.017.

32. A. A. Kolodziejczyk, J. K. Kim, J. C. Tsang, T. Ilicic, J. Henriksson, K. N. Natarajan, A. C. Tuck, X. Gao, M. Buhler, P. Liu, J. C. Marioni, and S. A. Teichmann. Single cell rna-sequencing of pluripotent states unlocks modular transcriptional variation. Cell Stem Cell, 17(4):471–85, 2015. ISSN 1875-9777 (Electronic) 1875-9777 (Linking). doi: 10.1016/j.stem.2015.09.011.

33. K. S. Song, P. E. Scherer, Z. Tang, T. Okamoto, S. Li, M. Chafel, C. Chu, D. S. Kohtz, and M. P. Lisanti. Expression of caveolin-3 in skeletal, cardiac, and smooth muscle cells. caveolin-3 is a component of the sarcolemma and co-fractionates with dystrophin and dystrophin-associated glycoproteins. J Biol Chem, 271(25):15160–5, 1996. ISSN 0021-9258 (Print) 0021-9258 (Linking). doi: 10.1074/jbc.271.25.15160.

34. A. Gurney, F. Axelrod, C. J. Bond, J. Cain, C. Chartier, L. Donigan, M. Fischer, A. Chaudhari, M. Ji, A. M. Kapoun, A. Lam, S. Lazetic, S. Ma, S. Mitra, I. K. Park, K. Pickell, A. Sato, S. Satyal, M. Stroud, H. Tran, W. C. Yen, J. Lewicki, and T. Hoey. Wnt pathway inhibition via the targeting of frizzled receptors results in decreased growth and tumorigenicity of human tumors. Proc Natl Acad Sci U S A, 109(29):11717–22, 2012. ISSN 1091-6490 (Electronic) 0027-8424 (Linking). doi: 10.1073/pnas.1120068109.

35. S. Yanagawa, F. van Leeuwen, A. Wodarz, J. Klingensmith, and R. Nusse. The dishevelled protein is modified by wingless signaling in drosophila. Genes Dev, 9(9):1087–97, 1995. ISSN 0890-9369 (Print) 0890-9369 (Linking). doi: 10.1101/gad.9.9.1087.

36. Nicole A. Repina, Xiaoping Bao, Joshua A. Zimmermann, David A. Joy, Ravi S. Kane, and David V. Schaffer. Optogenetic control of wnt signaling for modeling early embryogenic patterning with human pluripotent stem cells. bioRxiv, 2019. doi: 10.1101/665695.

37. S. J. Habib, B. C. Chen, F. C. Tsai, K. Anastassiadis, T. Meyer, E. Betzig, and R. Nusse. A localized wnt signal orients asymmetric stem cell division in vitro. Science, 339(6126):1445–8, 2013. ISSN 1095-9203 (Electronic) 0036-8075 (Linking). doi: 10.1126/science.1231077.

38. T. Schwarz-Romond, C. Merrifield, B. J. Nichols, and M. Bienz. The wnt signalling effector dishevelled forms dynamic protein assemblies rather than stable associations with cytoplasmic vesicles. J Cell Sci, 118(Pt 22):5269–77, 2005. ISSN 0021-9533 (Print) 0021-9533 (Linking). doi: 10.1242/jcs.02646.

39. M. Gagliardi, A. Hernandez, I. J. McGough, and J. P. Vincent. Inhibitors of endocytosis prevent wnt/wingless signalling by reducing the level of basal beta-catenin/armadillo. J Cell Sci, 127(Pt 22):4918–26, 2014. ISSN 1477-9137 (Electronic) 0021-9533 (Linking). doi: 10.1242/jcs.155424.

40. S. M. Jones, K. E. Howell, J. R. Henley, H. Cao, and M. A. McNiven. Role of dynamin in the formation of transport vesicles from the trans-golgi network. Science, 279(5350):573–7, 1998. ISSN 0036-8075 (Print) 0036-8075 (Linking). doi: 10.1126/science.279.5350.573.

41. G. Kreitzer, A. Marmorstein, P. Okamoto, R. Vallee, and E. Rodriguez-Boulan. Kinesin and dynamin are required for post-golgi transport of a plasma-membrane protein. Nat Cell Biol, 2(2):125–7, 2000. ISSN 1465-7392 (Print) 1465-7392 (Linking). doi: 10.1038/35000081.

42. M. Vinyoles, B. Del Valle-Perez, J. Curto, R. Vinas-Castells, L. Alba-Castellon, A. Garcia de Herreros, and M. Dunach. Multivesicular gsk3 sequestration upon wnt signaling is controlled by p120-catenin/cadherin interaction with lrp5/6. Mol Cell, 53(3):444–57, 2014. ISSN 1097-4164 (Electronic) 1097-2765 (Linking). doi: 10.1016/j.molcel.2013.12.010.

43. A. I. Hagemann, J. Kurz, S. Kauffeld, Q. Chen, P. M. Reeves, S. Weber, S. Schindler, G. Davidson, T. Kirchhausen, and S. Scholpp. In vivo analysis of formation and endocytosis of the wnt/beta-catenin signaling complex in zebrafish embryos. J Cell Sci, 127(Pt 18): 3970–82, 2014. ISSN 1477-9137 (Electronic) 0021-9533 (Linking). doi: 10.1242/jcs.148767.

44. V. Bryja, L. Cajanek, A. Grahn, and G. Schulte. Inhibition of endocytosis blocks wnt signalling to beta-catenin by promoting dishevelled degradation. Acta Physiol (Oxf), 190(1): 55–61, 2007. ISSN 1748-1708 (Print) 1748-1708 (Linking). doi: 10.1111/j.1365-201X.2007.01688.x.

45. S. Grainger, N. Nguyen, J. Richter, J. Setayesh, B. Lonquich, C. H. Oon, J. M. Wozniak, R. Barahona, C. N. Kamei, J. Houston, M. Carrillo-Terrazas, I. A. Drummond, D. Gonzalez, K. Willert, and D. Traver. Egfr is required for wnt9a-fzd9b signalling specificity in haematopoietic stem cells. Nat Cell Biol, 21(6):721–730, 2019. ISSN 1476-4679 (Electronic) 1465-7392 (Linking). doi: 10.1038/s41556-019-0330-5.

46. K. Labun, T. G. Montague, J. A. Gagnon, S. B. Thyme, and E. Valen. Chopchop v2: a web tool for the next generation of crispr genome engineering. Nucleic Acids Res, 44 (W1):W272–6, 2016. ISSN 1362-4962 (Electronic) 0305-1048 (Linking). doi: 10.1093/nar/gkw398.

47. E. K. Brinkman, A. N. Kousholt, T. Harmsen, C. Leemans, T. Chen, J. Jonkers, and B. van Steensel. Easy quantification of template-directed crispr/cas9 editing. Nucleic Acids Res, 46 (10):e58, 2018. ISSN 1362-4962 (Electronic) 0305-1048 (Linking). doi: 10.1093/nar/gky164.

